# Endothelial and non-endothelial responses to estrogen excess during development lead to vascular malformations

**DOI:** 10.1101/320234

**Authors:** Silvia Parajes, Sophie Ramas, Didier Y.R. Stainier

## Abstract

Excess estrogen signaling is associated with vascular malformations and pathologic angiogenesis, as well as tumor progression and metastasis. Yet, how dysregulated estrogen signaling impacts vascular morphogenesis *in vivo* remains elusive. Here we use live imaging of zebrafish embryos to determine the effects of excess estrogen signaling on the developing vasculature. We find that excess estrogens during development induce intersegmental vessel defects, endothelial cell-cell disconnections, and a shortening of the circulatory loop due to arterial-venous segregation defects. Whole-mount *in situ* hybridization and qPCR analyses reveal that excess estrogens negatively regulate Sonic hedgehog (Hh)/Vegf/Notch signaling. Activation of Hh signaling with SAG partially rescues the estrogen-induced vascular defects. Similarly, increased *vegfaa* bioavailability, using *flt1/vegfr1* mutants or embryos overexpressing *vegfaa_165_*, also partially rescues the estrogen-induced vascular defects. We further find that excess estrogens promote aberrant endothelial cell (EC) migration, possibly as a result of increased PI3K and Rho GTPase signaling. Using estrogen receptor mutants and pharmacological studies, we show that Esr1 and the G-protein coupled estrogen receptor (Gper1) are the main receptors driving the estrogen-induced vascular defects. Mosaic overexpression of *gper1* in ECs promotes vascular disconnections and aberrant migration, whereas no overt vascular defects were observed in mosaic embryos overexpressing wild-type or constitutively active nuclear estrogen receptors in their ECs. In summary, developmental estrogen excess leads to a mispatterning of the forming vasculature. Gper1 can act cell-autonomously in ECs to cause disconnections and aberrant migration, whilst Esr signaling predominantly downregulates Hh/Vegf/Notch signaling leading to impaired angiogenesis and defective arterial-venous segregation.

**Subject codes**: angiogenesis, animal models of human disease, mechanisms, vascular biology.

## INTRODUCTION

Vascular development is largely conserved across species, and studies in fish, birds and mammals have brought significant understanding into the main molecular mechanisms orchestrating vascular morphogenesis ^1^. Notochord-derived sonic hedgehog (HH) induces vascular endothelial growth factor A (*Vegfa*) expression in the ventral somites. VEGFA, via activation of kinase insert domain receptor (KDR/VEGFR2), orchestrates angioblast differentiation and proliferation, as well as vasculogenesis and angiogenesis during development and disease ^1,2^. VEGFA-dependent activation of phosphatidylinositol 3-kinase (PI3K)/AKT and the mitogen-activated kinase (MAPK) pathways activates the Rho family of small GTPases Rac1, Cdc42 and RhoA, to promote directional migration ^3,4^. Furthermore, VEGF signaling promotes arterial specification by inducing endothelial *Dll4* expression and activation of Notch signaling ^1,5^. Studies in zebrafish have shown that expression of the arterial and venous markers EphrinB2 (*efnb2a*) and EphB4 (*ephb4a*), respectively, in the dorsal aorta (DA) primordium is required for controlled ventral sprouting and arterial-venous segregation ^6^. Importantly, dysregulation of VEGFA signaling and downstream effectors leads to vascular defects, hence, hampering adequate nutrient and oxygen supply to irrigated tissues. *Vegfa* haploinsuficiency impairs angiogenesis and vasculogenesis and is embryologic lethal in mice. Furthermore, reduced Notch signaling induces arterial-venous malformations and promotes endothelial hypersprouting during angiogenesis ^1,5^. Hence, tight regulation of these signaling pathways is essential for the formation of a functional vascular system.

Estrogens are sex steroid hormones promoting capillary formation and angiogenesis ^7–9^. 17β-estradiol (E2), the most potent endogenous estrogen, exerts its biological action via two nuclear estrogen receptors ESR1 (ERα) and ESR2 (ERβ), and the G protein-coupled estrogen receptor GPER1. Classical estrogen signaling involves translocation of ESR1 and ESR2 into the nucleus, where they modulate gene transcription by directly binding to estrogen responsive elements (ERE) present in the promoter of target genes, or acting as co-factors of other transcription factors ^10^. Physiological nuclear ESR1 signaling induces transcriptional activation of *VEGFA* and *VEGFR2*^7,11^. In addition, non-genomic estrogen signaling promotes EC and progenitor EC (PEC) migration *in vitro* by inducing nitric oxide (NO) synthesis ^12^, synergizing with tyrosine kinase receptor signaling, or via interaction with PI3K or MAPK pathways and RhoA activation ^8,13–15^.

The significance of estrogen signaling in vascular morphogenesis is further supported by studies using genetic models of estrogen deficiency showing deficient ischemia reperfusion in *Esr1* knockout mice ^16–20^. On the other hand, increased estrogen levels in patients with liver disease are associated with a higher prevalence of angiomas ^21^. Furthermore, increased fetal lethality and placental hemorrhage has been reported in two genetic models of excess estrogen in mouse ^22,23^. In both cases, embryonic lethality was preceded by an increase in estrogen levels. Similar findings were reported when injecting E2 in pregnant mice ^22^. In addition to genetic conditions leading to estrogen excess, a large number of endocrine disruptors with estrogenic activity are present in the environment ^24^. However, studies into the molecular and cellular responses to excess estrogen signaling in the forming vasculature and its role in the etiology of pathologic angiogenesis, are still scarce.

Here, combining single-cell resolution microscopy together with genetic and pharmacologic studies, we found that excess estrogen induces a mispatterning of the forming vasculature. Vascular defects were a result of antiangiogenic cues due to a negative regulation of the Hh/Vegf/Notch signaling pathways, and exacerbated EC migration due to synergisms with PI3K and Rho GTPases signaling pathways. The E2-induced vascular malformations were more prominently mediated by Esr1 and Gper1. Finally, we also found that Gper1 can act cell-autonomously to induce EC migration and cell-cell disconnections, whilst nuclear estrogen signaling appears to act in non-endothelial cells. Altogether, our findings are of broad translational significance in endocrine and vascular diseases. Importantly, this study also reveals complex autocrine and paracrine responses to excess estrogen action cooperating in the developmental programming of vascular disease.

## MATERIAL AND METHODS

### 1. Transgenic zebrafish lines and animal husbandry

All zebrafish husbandry was performed under standard conditions in accordance with institutional (MPG) and national ethical and animal welfare guidelines. Wild-type AB zebrafish and the previously established zebrafish lines *Tg(kdrl:NLS-mCherry)^is4^* ^25^, *TgBAC(cdh5:GAL4FF)^mu101^* ^26^, *Tg(UAS:LIFEACT-GFP)^mu271^* ^27^, *Tg(kdrl:EGFP)^s843^* ^28^, *TgBAC(etv2:EGFP)^ci1^* ^29^, *Tg(hsp70l:vegfaa_165_,cryaa:cerulean)^s712^* ^30^, *Tg(5xERE:GFP)^c262^* ^31^, *kdrl^hu5088^* ^32^*, flt1^bns29^* ^33^, and *vegfaa^bns1^* ^34^ were used in this study. To improve readability, *TgBAC(cdh5:GAL4FF); Tg(UAS:LIFEACT-GFP)* was simplified to *LIFEACT-GFP*, and *Tg(kdrl:NLS-mCherry)* is referred to as *NLS-mCherry*. Zebrafish embryos were obtained from natural spawning of the aforementioned zebrafish lines and raised at 28 C in egg water.

### 2. Chemical treatments

Chemical treatments of zebrafish embryos were conducted as described in supplementary methods.

### 3. Gene expression analyses

Twenty vehicle or E2-treated embryos were collected at 30 hpf in 400 µL of Trizol^®^ (Thermo Fisher Scientific, Schwerte, Germany) for total RNA extraction. RT-PCR was performed using 3 µg of RNA and the Maxima First Strand cDNA Synthesis Kit. Synthesized cDNA was DNaseI treated (Thermo Fisher Scientific). qPCR was performed using DyNAmo ColorFlash SYBR Green (Thermo Fisher Scientific), 0.5 µL cDNA and 300 nM primers in a CFX Connect^TM^ Real-Time System (Bio-Rad GmbH, München, Germany). Expression studies were conducted in at least 3 biological replicates. *rpl13* and *elfα* were used as housekeeping genes. Gene expression was normalized to control using the 2^-∆∆Ct^ method. Graphs were generated using Prism version 6.0 (Graphpad software Inc.).

Expression of *vegfaa, shha, and ptch1* was analyzed by whole-mount in situ hybridization (WISH) as previously described ^35^. Vehicle or E2-treated embryos were collected at 30 hpf and fixed overnight in fish fixative containing 4% paraformaldehyde, 22.6 mM NaH_2_PO_4_, 77 mM Na_2_HPO_4_, 1.2 mM CaCl_2_, 40% Sucrose (all chemicals were purchased from Sigma-Aldrich). Digoxigenin (DIG)-labeled cRNA probes were synthesized using a DIG RNA labeling Kit (Roche, Berlin, Germany). Primers are listed in Suppl. Table 1. Brightfield images were acquired with a SMZ25 stereomicroscope (Nikon).

### 4. Assessment of vascular defects

Vascular defects were evaluated in control and treated *Tg(kdrl:EGFP)* or *TgBAC(etv2:EGFP*) embryos under a fluorescent stereo microscope (Zeiss) at 48 hpf (days post-fertilization). The number of missing or disconnected intersegmental vessels (ISVs) on each side of the 9 somites anterior to the cloaca was quantified (i.e. 20 ISVs). The length of the lumenized DA reflects the number of somites with a lumenized DA. Data were obtained from at least 5 embryos from 3 biological replicates. Graphs were generated using Prism.

### 5. Imaging

*LIFEACT-GFP* or *LIFEACT-GFP; NLS-mCherry* embryos were used for time-lapse confocal imaging of the DA, ISVs and the common cardinal vein (CCV). Embryos were mounted in a glass-bottom petri dish (MatTek), using 0.6% low melting agarose containing 0.08 mg/mL tricaine (MS-222, Sigma-Aldrich), and 8 µM E2 or vehicle. The petri dish was filled with egg water containing 0.04 mg/mL tricaine and 8 µM E2 or vehicle. Time-lapse imaging was performed under controlled heating conditions adjusted at 28.5 C. Images of the forming ISVs and the most posterior region of the DA were acquired every 20 min using a LD C-apochromat 40X/1.1 W objective lense in a LSM 800 inverted (Axio Observer) confocal laser scanning microscope (Zeiss). Images of the forming CCV were acquired every 15 min using the same settings in a LSM 800 inverted confocal microscope or a Spinning disk inverted CSU-X1 microscope (Zeiss).

*Tg(kdrl:EGFP)* embryos were used for confocal imaging of the vascular defects. Embryos were collected at 48 hpf and fixed overnight in fish fixative. Fixed embryos were mounted in 1% low melting agarose. Z-stack confocal images were acquired using a LSM 800 inverted confocal microscope as described above. Brightfield images of anaesthetized 48 hpf embryos were acquired with a SMZ 25 stereomicroscope (Nikon).

### 6. Surface rendering

Confocal images of *LIFEACT-GFP; NLS-mCherry* embryos were used for surface rendering of LIFEACT-GFP and nuclear mCherry expression using automatic surface detection in Imaris x64 version 8.4.1 (Bitplane AG). Cortical endothelial LIFEACT-GFP expression was used for manual surface rendering of individual using Imaris.

### 7. Migration analyses

Time-lapse images of the forming CCV in *LIFEACT-GFP; NLS-mCherry* embryos were used for migration analyses. EC mCherry+ were manually selected using spot detector, and tracked over time using spot tracker in Imaris. At least 7 ECs were tracked for each sample from 3 biological replicates. Data on migration track length, migration track displacement length and migration persistence (migration track length/migration length) were obtained using Imaris track analyses, and plotted using Prism.

### 8. Generation of nuclear estrogen receptor mutants

Mutant *esr1^bns229^*, *esr2a^bns228^* and *esr2b^bns230^* alleles were generated using the CRISPR/Cas9 system. The *esr1^bns229^* allele contains a 7 bp deletion (c.477delGCAGCCG; p.A160Wfs*97) resulting in a predicted truncated polypeptide containing 160 aa. The *esr2a^bns228^* allele carries a 4 bp deletion (c.427delCAGA; p.T143Rfs*6) and encodes for a predicted polypeptide containing 143 aa. The *esr2b^bns230^* allele carries a 1 bp deletion and an insertion of 9 bp (c.441del1ins9; p.D147Gfs*6), which encodes for a predicted polypeptide containing 147 aa of the N-terminus of the wild-type protein. Small guide RNAs (sgRNA) were designed against exon 3 of each gene, which contains the DNA binding domain. Single stranded oligos containing the sense and antisense sequence of the sgRNA (Supplemental Table 1) were annealed and cloned into the linearized pT7-gRNA vector. Cloned sgRNA vectors were linearized with BamHI and used to synthesize the sgRNA with the MEGAshortscript T7 kit (Ambion, Kaufungen, Germany). The Cas9 vector (pT3TS-nlsCas9nls, Addgene) was linearized with XbaI and Cas9 mRNA was synthesized using the mMESSAGE mMACHINE T3 transcription kit (Ambion). Synthesized RNA was purified using the RNA Clean & Concentrator-5 kit (Zymo Research, Freiburg, Germany). One nL of an injection solution containing 6.5 ng/µL sgRNA and 150 ng/µL *Cas9* mRNA was injected into 1-cell stage embryos.

### 9. Genotyping of mutant alleles

*kdrl^hu5088^*, *esr1^bns229^*, *esr2a^bns228^* and *esr2b^bns230^* animals were genotyped by high-resolution melting curve analyses using DyNAmo ColorFlash SYBR Green (Thermo Fisher Scientific) and specific primers (Supplemental Table 1) in an Eco Real-Time PCR system (Illumina). Genotypes were analyzed on normalized derivative plots. *flt1^bns29^* embryos were genotyped as previously described ^33^.

### 10. Mosaic endothelial overexpression of estrogen receptors

The coding sequence of *esr1, esr2a, esr2b*, or *gper1* was cloned downstream the P2A signal in a *fli1ep:membrane-tdTomato-P2A* using the ClaI restriction site in a pT2 backbone containing tol2 transposon terminal sequences. A constitutively active *esr1* mutant carrying the p.Y549S missense mutation was generated by site-directed mutagenesis using primers listed on Supplemental Table 1.

One-cell stage *Tg(kdrl:EGFP)* or *Tg(ERE:GFP)* embryos were injected with 30 pg of the generated constructs and 25 pg *tol2* mRNA. Live embryos showing membrane tdTomato (mTomato) expression were imaged using LSM 800 inverted confocal laser scanning microscope at 24 hpf as described above.

### 11. Statistical analyses

Data following a normal distribution were analyzed using a t-test or a one-way analysis of variance (ANOVA) with Tukey correction for multiple comparisons. Data not showing a Gaussian distribution were compared using the non-parametric Mann-Whitney test or a Kruskal-Wallis test with Dunn’s correction for multiple comparisons. Statistical analyses were performed using Prism.

## RESULTS

### Excess estrogens impact angiogenesis and vasculogenesis

No overt developmental abnormalities were observed in E2-treated zebrafish embryos at 48 hpf (Figure 1A-A’). However, analysis of *Tg(kdrl:EGFP)* embryos revealed that excess estrogens induced vascular defects including missing or disconnected ISVs, ectopic sprouting from the dorsal longitudinal anastomotic vessel (DLAV), and ISV stenoses (Figure 1B’). The ISV defects were dose-dependent. Non-linear regression analyses (R square 0.8860) calculated an EC50 of 4 µM, with the induced phenotypes being significantly different from controls starting at 1 µM (Figure 1C).

**Figure 1.**
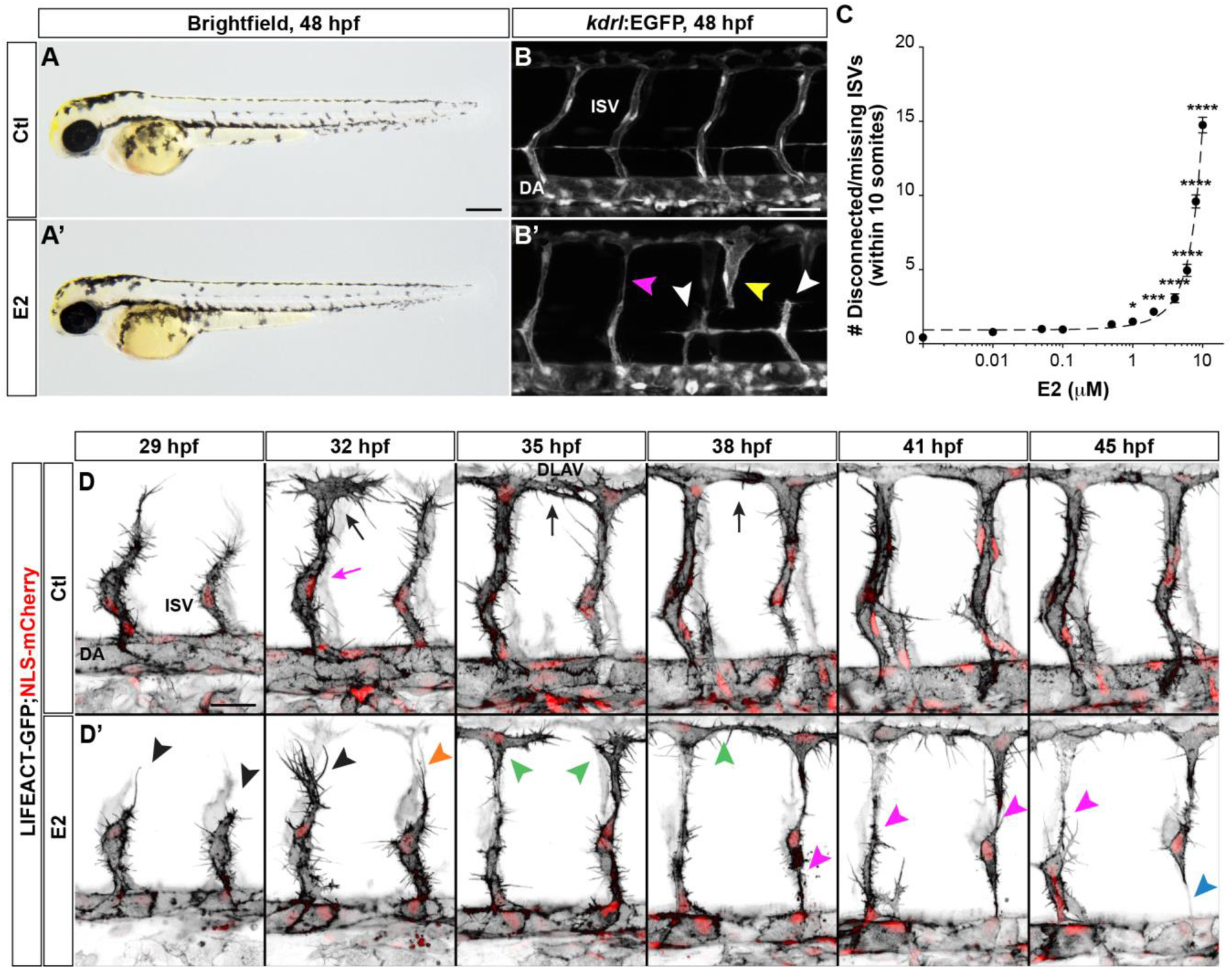
Excess estrogens during developmental angiogenesis induce EC cell-cell disconnections and ISV defects. (**A-A’**) Brightfield images of 48 hpf embryos treated with vehicle (Ctl) or E2. E2 treatment induce no overt morphological defects (n=10). (**B-B’**) Maximum intensity projections of the trunk vasculature anterior to the cloaca of Ctl (B) or E2-treated (B’) embryos at 48 hpf. E2-treated embryos exhibit stenosed (pink arrowhead) and missing/disconnected (white arrowheads) intersegmental vessels (ISVs), and ectopic sprouting from the dorsal longitudinal anastomotic vessel (DLAV, yellow arrowhead). **(C)** E2 dose-response curve for the ISV phenotype at 48 hpf. X-axis is in logarithmic scale. The E2-induced ISV defects become significantly different from control embryos at 1 µM (n=20). Mean ± SEM are shown. (**D-D’**) Time-lapse confocal imaging of forming ISVs in Ctl (D) and E2-treated (D’) embryos between 29-45 hpf. Negative maximum intensity projections of endothelial actin filaments (LIFEACT-GFP, black) and nuclei (NLS-mCherry, red) are shown. Control ISVs migrate dorsally, undergo lumen formation (pink arrow) and anastomose to form the dorsal longitudinal anastomotic vessel (DLAV) (black arrows). ISVs in E2-treated embryos appear delayed (black arrowheads), yet extend dorsally and form the DLAV (green arrowheads). During lumen formation, ISVs show stenoses (pink arrowheads) and disconnect from the dorsal aorta (DA) (blue arrowhead). *, p<0.05; ***, p<0.001; ****, p<0.0001. Scale bars, 250 µm (A-A’); 50 µm (B-B’) and 25 µm (D-D’).

We next performed time-lapse confocal imaging during ISV formation in the posterior region of the trunk of *LIFEACT-GFP;NLS-mCherry* embryos, reporting for endothelial filamentous actin (F-actin) and nuclei, respectively, between 29-44 hpf. In control embryos, ECs in the forming ISVs migrated dorsally from the DA to then anastomose and form the DLAV. By 35 hpf, this process was completed and ISV lumen formation and cardinal vein (CV) sprouting were also observed (Figure 1D and Movie 1A). Similarly to controls, tip ECs in E2-treated embryos projected actin rich filopodia during dorsal migration and anastomosed to form the DLAV. However, dorsal EC migration in E2-treated embryos was delayed and stalling at the midline was observed in some ISVs (Figure 1D’ and Movies 1B-C). Although hampered, these projections eventually connected to ECs in the DLAV (Movie 1B). Strikingly, strong narrowing of the forming ISVs was observed from 38 hpf, coinciding with lumen formation. These stenoses resolved with loss of cell-cell contacts and ISVs disconnecting from the DLAV or DA (Figure 1D’ and Movies 1B-C).

In addition to the ISV defects, 48 hpf E2-treated embryos had a shorter circulatory loop due to a premature truncation of the lumenized DA (Figure 2A-A’). At 24 hpf, when treatments were initiated, arterial-venous segregation and extension of the circulatory loop are not yet completed. Time-lapse imaging in control *LIFEACT-GFP* embryos showed active expansion of the DA lumen between 28-43 hpf (Movie 2A). Transient stenoses and cell shape changes were observed within regions of lumen expansion (Figure 2C and Movie 2A). In E2-treated embryos, however, expansion of the lumenized DA was stalled. Stenoses were also observed at presumptive lumen extension regions. In contrast to controls, stenoses did not resolve in E2-treated embryos. Strong cortical F-actin expression was observed in ECs. Furthermore, cells at the most posterior end of the DA became elongated and migrated posteriorly and ventrally to exit the lumenized DA. Actively migrating ECs lost anterior connections with neighboring ECs, resulting in a premature truncation of the lumenized DA and, hence, the circulatory loop (Figure 2C’ and Movie 2B). The DA phenotype was also dose-dependent (Figure 2B). E2-induced the DA phenotype starting at 2 µM, and non-linear regression analyses (R square 0.9216) calculated an EC50 of 5 µM.

**Figure 2.**
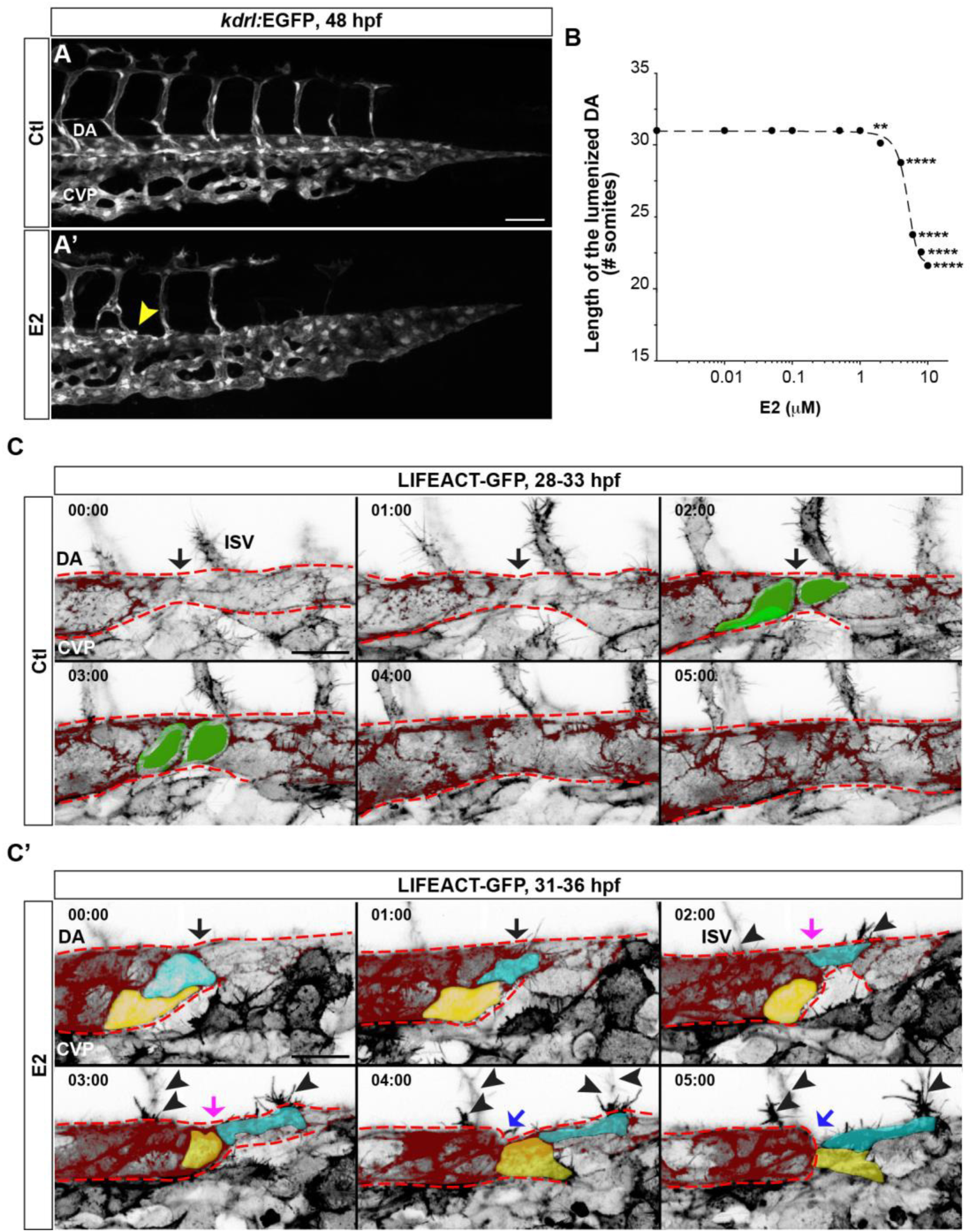
Excess estrogens induce a shortening of the circulatory loop due to impaired axial vessel segregation. **(A)** Maximum intensity projections of the posterior trunk vasculature of vehicle (Ctl) (A) or E2-treated (A’) embryos at 48 hpf. Excess estrogens induce a premature truncation of the lumenized dorsal aorta (DA) (yellow arrowhead). **(B)** E2 dose-response curve for the DA phenotype at 48 hpf. X-axis is in logarithmic scale. E2 effects on the DA becomes significantly different at 2 µM (n=20). Mean ± SEM are shown. (**C-C’**) Time-lapse confocal images of the DA lumen expansion of Ctl (C) and E2-treated (C’) embryos. Negative images of endothelial F-actin (LIFEACT-GFP) are shown. Time is at the top left corner (hh:mm). The DA is pseudocolored in red and outlined with red dashed lines. DA stenoses are observed within the lumen expansion region (black arrows) (C-C’). (C) Rendered in green, control ECs undergoing transient cell shape changes during lumen expansion. (C’) DA stenoses in E2-treated embryos fail to resolve (pink arrow). Rendered in yellow and blue, two ECs at the stenosed region migrating posteriorly and detaching from their anterior neighbors; resulting in a premature truncation of the circulatory loop (blue arrows). Black arrowheads, delayed intersegmental vessel (ISV) sprouting. *, p<0.05; ****, p<0.0001. Scale bars, 50 µm (A) and 20 µm (C-C’). CVP, cardinal vein plexus.

Treatments with A4 and T, two E2 precursors, (Supplemental Figure 1A) also induced ISV disconnections and a premature truncation of the lumenized DA by 48 hpf (Supplemental Figure 1B-B’). Co-incubation with letrozole, an aromatase inhibitor blocking their conversion into E2, partially rescued the ISV and DA defects in A4. T induced milder ISV defects and a partly penetrant DA phenotype (5/39 embryos). Although not statistically significant, letrozole also improved the vascular defects in T-treated embryos (Supplemental Figure 1B-B’). A4 and T can also be metabolized into 11-KT and DHT, two potent androgens (Supplemental Figure 1A). Treatments with 11KT and DHT had no effects on vascular morphogenesis (Supplemental Figure 1C-C’). Altogether, these findings indicate that the described vascular malformations are estrogen-specific.

### Excess estrogens negatively regulate the Hh/Vegf/Notch signaling pathways during vascular development

The E2-induced ISV phenotype was indicative of angiogenic defects. Therefore, we first assessed the expression of Vegf ligand and receptor genes, master regulators of developmental angiogenesis. qPCR analyses performed at 30 hpf, just before the onset of the vascular defects, revealed a 48% reduction in *vegfaa* expression compared to controls (Figure 3A). Reduced *vegfaa* expression in the posterior somites of E2-treated embryos was observed by WISH (Figure 3A’). No changes in the expression of other Vegf ligand genes were observed. Similarly, expression of *kdrl/vegfr2, flt4/vegfr3*, and the decoy *vegfaa* receptor ^1^, *flt1/vegfr1*, appeared unchanged (Figure 3A).

**Figure 3.**
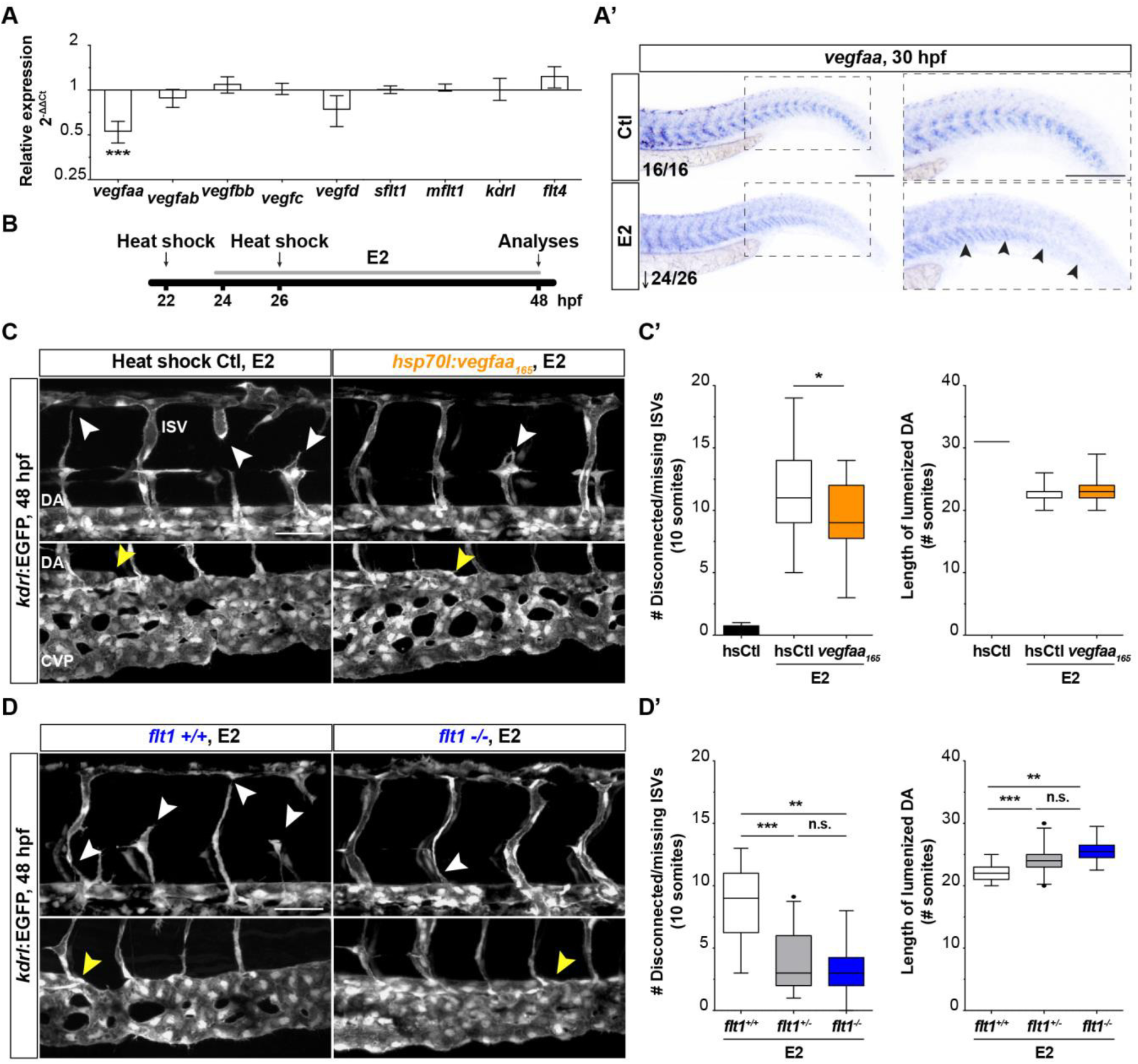
Developmental estrogen excess negatively regulates Vegfaa signaling. **(A-A’)** Expression analyses of *vegf* ligand and receptor genes in 30 hpf vehicle (Ctl) and E2-treated embryos using qPCR (A) and whole-mount *in situ* hybridization (A’). Excess estrogens downregulate somitic *vegfaa* expression (black arrowheads). Mean±SEM are plotted. **(B)** Schematic of the heat shock (hs) protocol used to rescue *vegfaa* expression in E2-treated embryos. **(C-C’)** Maximum intensity projections (C) and quantification (C’) of the E2-induced intersegmental vessel (ISV) and dorsal aorta (DA) phenotypes in heat shocked *(hsp70l:vegfaa165)* and wild-type siblings at 48 hpf. *vegfaa_165_* overexpression partially rescues the ISV (white arrowheads), and, albeit not significant, the DA phenotypes (yellow arrowheads). **(D-D’)** Maximum intensity projections (D) and quantification (D’) of the E2-induced vascular defects in *flt1* mutants and wild-type siblings. Increased Vegfaa bioavailability in *flt1* mutants significantly rescues the E2-induced ISV (white arrowheads) and DA (yellow arrowheads) phenotypes. Top panels in (C, D), trunk vasculature anterior to the cloaca. Bottom panels in (C, D), axial vessels within somites 22 and 26. *, p<0.05, **, p<0.01, ***, p<0.001; n.s., not significant. Scale bars, 200 µm (A’) and 50 µm (C, D). *sflt1*, soluble flt1; *mflt1*, membrane flt1; CVP, cardinal vein plexus; ↓, reduced *vegfaa* expression.

We next treated with E2 *Tg(hsp70l:vegfaa_165_)* embryos heat shocked at 22 and 26 hpf (Figure 3B). *vegfaa_165_* overexpression significantly reduced the number of disconnected or missing ISVs, compared to controls (9 vs 11; median) (Figure 3C-C’); and improved, albeit not in a statistically significant way, the DA phenotype (24 vs 23; median). Similarly to what was observed in the *vegfaa* overexpression studies, upon E2 treatment *flt1^-/-^* and *flt1^+/-^* embryos exhibited a significantly lower number of ISV defects than wild-type siblings (3 vs 3 vs 9; median) (Figure 3D-D’). Also, *flt1^-/-^* and *flt1^+/-^* embryos exhibited a longer lumenized DA, compared to wild-type siblings (24 vs 24 vs 22; median) (Figure 3D-D’). These results indicate that reduced *vegfaa* expression plays a role in the etiology of the E2-induced vascular defects.

Hh signaling induces *Vegfa* expression ^1,2^. No significant changes in *shha*, *smo*, and *gli3* expression were observed by qPCR and WISH at 30 hpf. However, E2 reduced expression of the Hh regulated genes *gli1* (27%), and *ptch1* (68%), and the latter nearly became undetectable in the myotome by WISH (Figure 4A-A’). The expression of *crlra*, a downstream target of Hh signaling, was mildly but not significantly reduced (Figure 4A). Activation of Hh signaling with the Smo agonist SAG induced a partial rescue of the E2-induced ISV phenotype, compared to DMSO controls (12 vs 14, median), and increased the length of the lumenized DA (23 vs 22, median) (Figure 4C-C’). These results indicate that excess estrogen signaling is a negative regulator of Hh signaling.

**Figure 4.**
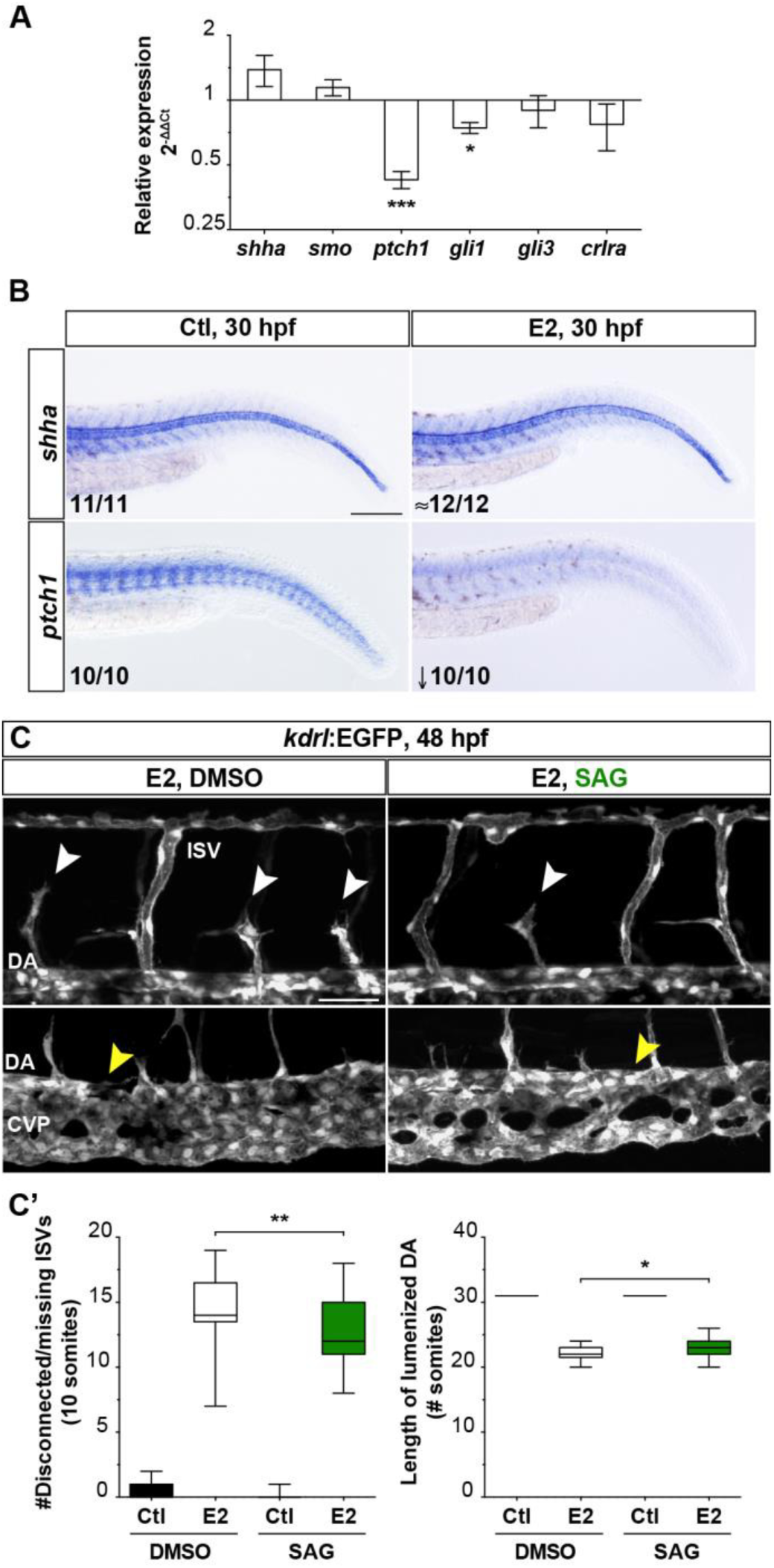
Excess estrogens impair Hh signaling. **(A-B)** Expression analyses of ligand and receptor genes of the Hh signaling pathway using qPCR (A) and whole-mount in situ hybridization (B) in 30 hpf control (Ctl) and E2-treated embryos. E2-treated embryos have reduced *gli1* expression and almost negligible myotome *ptch1* expression. Mean ± SEM are shown. **(C-C’)** Maximum intensity projections (C) and quantification (C’) of the effect of the smoothened agonist SAG on E2-induced vascular defects at 48 hpf. Hh signaling activation partially rescues the intersegmental vessel (ISV) (white arrowheads) and dorsal aorta (DA, yellow arrowheads) phenotypes induced by excess estrogens. Top panels in (C), trunk vasculature anterior to the cloaca. Bottom panels in (C), axial vessels within somites 22 and 26. *, p<0.05, **, p<0.01, ***, p<0.001. Scale bars, 200 µm (A) and 50 µm (C, D). CVP, cardinal vein plexus. ≈, unchanged *shha* expression; ↓, reduced *ptch1* expression.

VEGFA-dependent activation of Notch signaling is required for arterial-venous segregation. *vegfaa^+/-^* embryos exhibit a mild and lowly penetrant arterial-venous malformation phenotype (6/40 embryos) (Supplemental Figure 2A). Treatments of *vegfaa^+/-^* embryos with 1 µM E2, an E2 dose with no effects on the expansion of the circulatory loop, led to a significant shortening of the lumenized DA when compared to untreated *vegfaa^+/-^* embryos or E2-treated wild-type siblings (29 vs 31 vs 31, respectively; median) (Supplemental Figure 2A-A’). Reduced expression of the Notch ligand gene *dll4* (28%) was observed in E2-treated embryos by qPCR at 30 hpf. Expression of *notch1b* and *notch3*, and the Notch-regulated arterial maker genes *efnb2a* or *hey2* appeared unaffected (Supplemental Figure 2B). Similarly to what was observed in *vegfaa^+/-^* embryos, injection of a low dose of a splice *dll4* MO ^36^ (1.5 ng) led to a mild and partly penetrant shortening of the circulatory loop (20/50 embryos). E2 treatment (4 µM) significantly worsened the DA phenotype in *dll4* morphants, compared to untreated *dll4* MO or uninjected E2-treated embryos (19 vs 30 vs 24; median) (Supplemental Figure 2C-C’). *dll4* morphants also exhibited ectopic sprouting in the dorsal side of the intersomitic space by 48 hpf. E2 treatment did not affect the *dll4* MO-dependent ectopic sprouting. Furthermore, injection of the *dll4* MO had no effects on the ISV defects induced by excess estrogens (Supplemental Figure 2D-D’). Altogether, these data indicate that excess estrogens impair vascular development due to a negative regulation of the Hh/Vegf/Dll4 axis, with significant downregulation of *ptch1*, *vegfaa*, and *dll4* expression following E2 treatment.

### Excess estrogens promote EC migration independently of Vegfaa by modulating PI3K and Rho GTPase signaling

A reduced number of disconnected/missing ISVs and a longer lumenized DA were observed in E2-treated *kdrl^+/-^* embryos, compared to wild-type siblings (ISVs, 5 vs 9; DA, 24 vs 23; median) (Supplemental Figure 3), suggesting that E2 exposure might affect vascular development in a Vegfaa/Kdrl-independent manner.

To investigate potential Vegfaa-independent effects of excess estrogens during vascular development, we examined how E2 treatment impact CCV formation. The CCV forms from the cardinal vein (CV) and extends around each side of the yolk to converge cranioventrally at the sinus venosus of the heart. This process relies on EC proliferation and collective cell migration, which are dependent on Vegfc and Cadherin 5 (Cdh5) signaling respectively ^27^. Time-lapse imaging in *LIFEACT-GFP;NLS-mCherry* embryos revealed an increased length of the CCV in E2-treated embryos (Figure 5C-C’). Migration analyses of individual EC nuclei in the forming CCV between 32-36 hpf revealed a 30% increased EC migration track length in E2-treated embryos. A milder increase (12%) on EC track displacement length was also observed in E2-treated embryos, indicative of reduced directional migration (migration persistence) (Figure 5E). Increased migration track length was more prominent in follower ECs (data not shown), whilst reduced directionality was more common in leader ECs (Movies 3A-B). These findings show that excess estrogens can promote aberrant EC migration in a Vegfaa-independent context.

**Figure 5.**
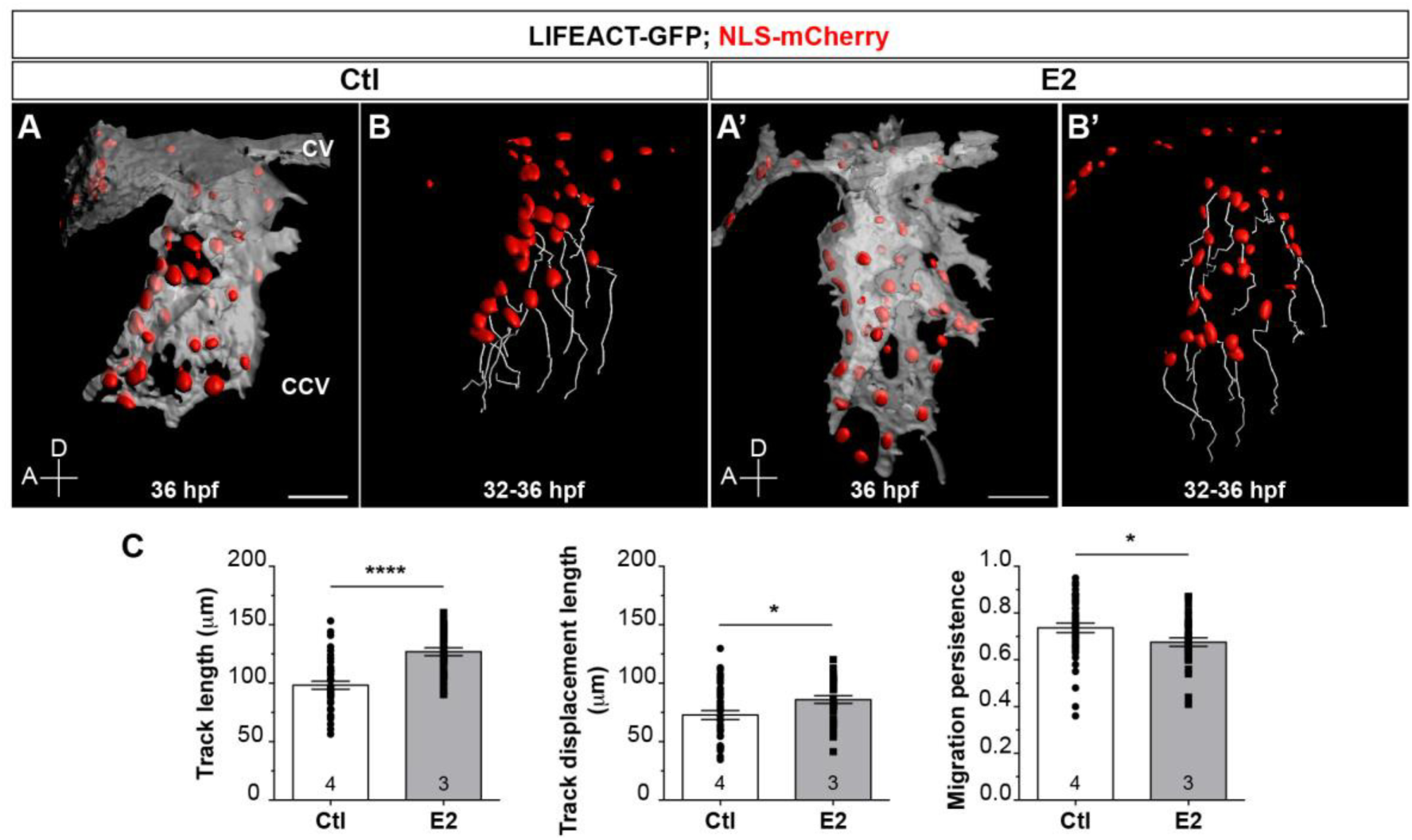
Excess estrogens promote EC migration in the common cardinal vein (CCV). **(A-C)** EC migration analyses in the forming CCV of Ctl (A-B) and E2-treated (A’-B’) embryos between 32-36 hpf. Three-dimensional rendering of endothelial LIFEACT-GFP (white) and nuclear NLS-mCherry (red) are shown. Migration paths (B-B’) and parameters of individual ECs were analyzed (C). E2 treatment significantly increase EC migration track length and displacement, but reduce directionality. Aligned scattered plots ± SEM are shown. Numbers in graphs indicate number of embryos analyzed. *, p<0.05; ****, p<0.0001. A, anterior; D, dorsal; CV, cardinal vein. Scale bars, 50 µm.

We next asked whether increased EC migration was contributing to the E2-induced vascular defects. First, we co-treated zebrafish embryos with E2 and the broad range PI3K inhibitor LY294002. The concentration of LY294002 used (1 µM) did not induce any vascular or morphological abnormalities (Figure 6A’). PI3K inhibition partially rescued the vascular defects induced by excess estrogens compared to DMSO controls (ISV, 12 vs 14; DA, 23 vs 22; median) (Figure 6A-A’). Similarly, inhibition of FAK, Rac1 and Rho induced a 40-50% reduction in E2-induced ISV defects. Although not statistically significant, fewer ISV defects were also observed upon Cdc42 inhibition (7 vs 8; median) (Figure 6B). Rac1, Cdc42 and Rho inhibition significantly increased the length of the DA in E2-treated embryos, compared to vehicle controls (24 vs 23, 25 vs 24, and 24 vs 23, respectively; median). FAK inhibition had no effects on the DA phenotype (Figure 6B’). These results indicate that increased PI3K and Rho GTPase signaling contributes to the E2-induced vascular defects.

**Figure 6.**
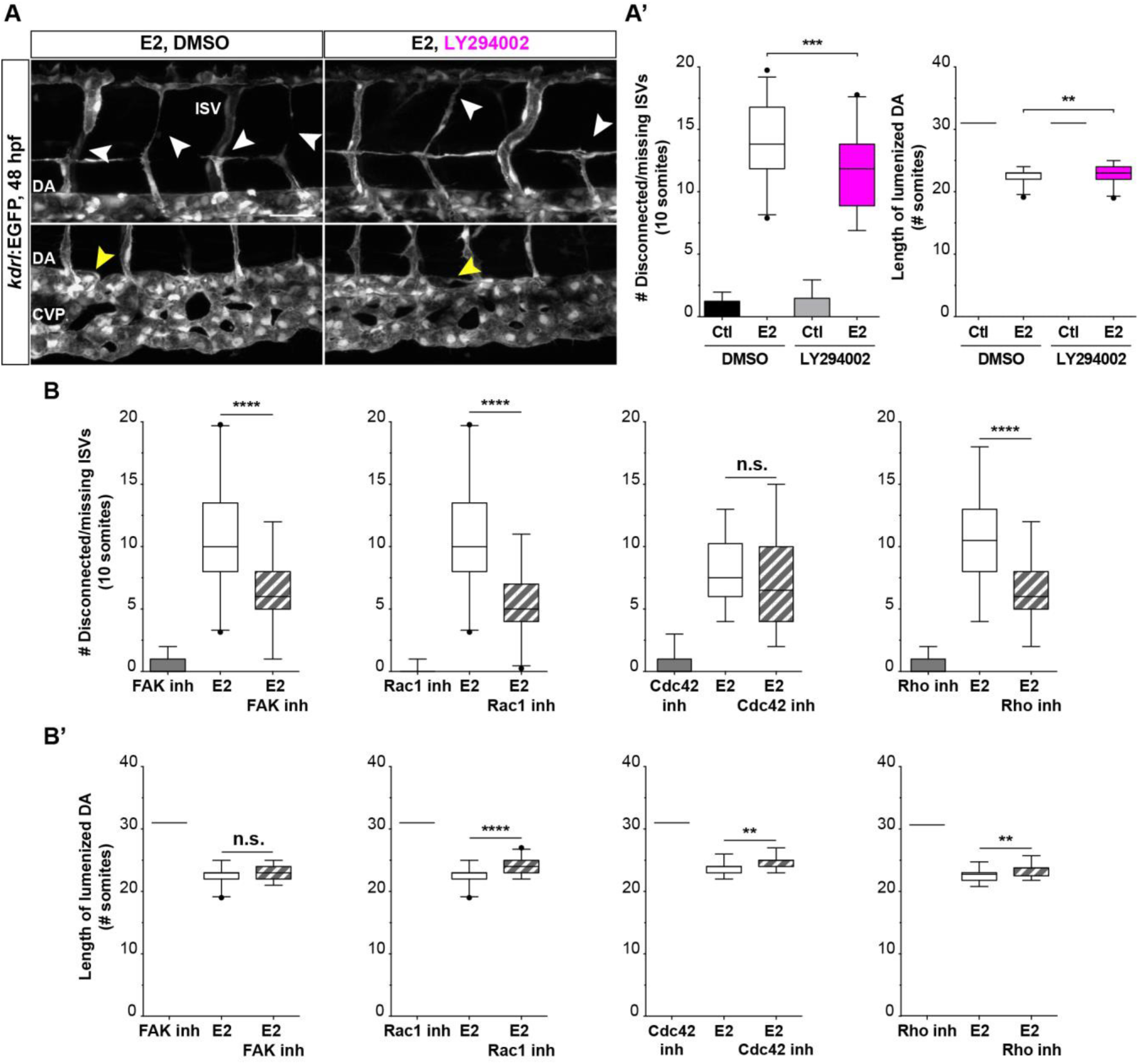
Increased PI3K and Rho GTPase signaling correlates with the vascular defects induced by excess estrogens. **(A-A’)** Maximum intensity projections (A) and quantification (A’) of the effect of the PI3K inhibitor LY29002 on the E2-induced vascular defects at 48 hpf. PI3K inhibition significantly rescues the intersegmental vessel (ISVs) (white arrowheads), and dorsal aorta (DA) phenotypes (yellow arrowheads) induced by excess estrogens. Top panels, trunk vasculature anterior to the cloaca. Bottom panels, axial vessels between somites 22 and 26. **(B-B’)** Quantification of the effect of focal adhesion kinase (FAK) and Rho GTPase inhibition on the E2-induced ISV (**B**) and DA (**B’**) phenotypes at 48 hpf. FAK, Rac1 and Rho inhibition significantly reduce the number of ISV defects in E2-treated embryos. Inhibition of Rac1, Cdc42 and Rho partially rescues the E2-induced DA phenotype. **, p<0.01; ***, p<0.001;****, p<0.0001; n.s., not significant. Scale bars, 50 µm. CVP, cardinal vein plexus.

### Signaling from Esr1 and Gper1 mediate the vascular defects induced by excess estrogens

Treatments with the pan-ER inhibitor ICI dramatically reduced the E2-induced ISV defects (2 vs 8; median) and increased the length of the lumenized DA (24 vs 22; median) (Figure 7A-B’). In zebrafish, the nuclear estrogen receptor α is encoded by *esr1*, whilst the nuclear estrogen receptor β is encoded by the paralog genes *esr2a* and *esr2b*. We generated mutant alleles for the three zebrafish nuclear receptor genes using the CRISPR/Cas9 system. The recovered *esr1^bns229^*, *esr2a^bns230^* and *esr2b^bns228^* alleles are predicted to encode proteins that lack the DNA binding, hinge, and ligand-binding domains and thus to be non-functional polypeptide (Supplemental Figure 4). *esr1*, *esr2a* and *esr2b* mutants have no overt morphological or vascular defects. E2 treatment in *esr1* mutants revealed a modest albeit significant rescue of the ISV and DA defects compared to wild-type siblings (ISV, 14 vs 16 and DA, 23 vs 22, respectively; median) (Figure 7C). *esr2a* and *esr2b* mutations had no significant effects on the E2-induced vascular defects induced by E2 (Figure 7C’-C’’).

**Figure 7.**
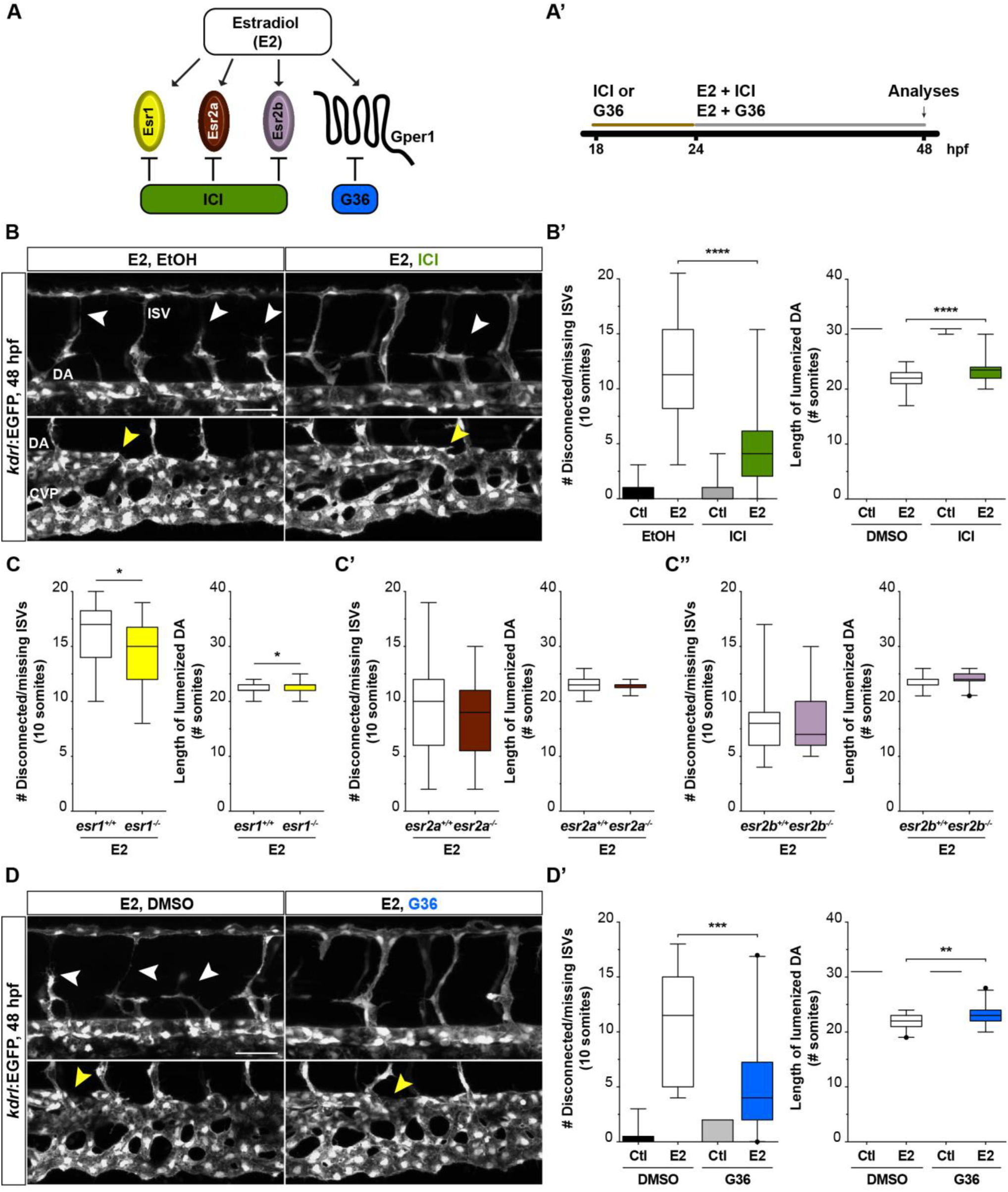
Esr1 receptor and Gper1 are the main drivers of the E2-induced vascular defects. (**A**) Schematic representation of the zebrafish estrogen receptors and inhibitors. **(A’)** Summary of the pharmacological approached used to block estrogen signaling. (**B-B’**) Maximum intensity projections (B) and quantification (B’) of the impact of the pan-estrogen receptor inhibitor, ICI, on the E2-induced vascular phenotypes. Inhibition of nuclear estrogen receptor signaling partially rescues the intersegmental vessel (ISV, white arrowheads) and dorsal aorta (DA, yellow arrowheads) defects induce by excess estrogens. **(C-C’’)** Quantification of the E2-induced vascular defects in 48 hpf *esr1*, *esr2a*, and *esr2b* mutants and wild-type siblings. Only *esr1^-/^*^-^ embryos are partially protected against the E2-induced vascular defects. **(D-D’)** Maximum intensity projections (D) and quantification (D’) of the effect of G36 on the E2-induced vascular phenotype. Selective Gper1 inhibition partially rescues the number ISV (white arrowheads) and DA (yellow arrowheads) phenotype induced by E2 treatment. Top panels in (B) and (D), trunk vasculature anterior to the cloaca. Bottom panels in (B) and (D), axial vessels between somites 22 and 26. *, p<0.05; **, p<0.01; ***, p<0.001; ****, p<0.0001. Scale bars, 50 µm. CVP, cardinal vein plexus.

In addition to the Esr receptors, estrogens also signal via Gper1. Selective Gper1 inhibition with G36 also induced a pronounced reduction of the ISV defects induced by E2 (4 vs 11.5; median) and increased the length of the circulatory loop (23 vs 22; median) (Figure 7D-D’). G1, a selective Gper1 agonist, also induced ISV disconnections that mimicked those induced by E2 (Supplemental Figure 5A). In addition, G1 treatments led to morphological defects in the most posterior region of the trunk vasculature. However, these defects did not recapitulate the DA defects induced by E2 (Supplemental Figure 5A). G1-induced vascular defects were fully rescued by G36 (Supplemental Figure 5A-A’). Hence, Esr1 and Gper1 are the main effectors of excess estrogens, although cooperative interactions with Esr2a and/or Esr2b might also contribute to the E2-induced vascular defects.

### Endothelial *gper1* signaling promotes migration and cell-cell disconnections

To ascertain whether the effects on EC migration and cell-cell contacts were due to a direct endothelial response to excess estrogens, we generated endothelial mosaic overexpression of the estrogen receptors. To enable the visualization of ECs overexpressing the estrogen receptors, constructs also contained *mTomato*. Control animals, injected with the construct *fli1ep:mTomato-P2A-EGFP*, exhibited no significant vascular defects at 24 hpf (Figure 8A-A’). Endothelial mTomato+ clusters were observed on the dorsal side of 24 hpf embryos injected with *fli1ep:mTomato-P2A-gper1* (6/8 embryos), and were disconnected from the forming ISVs (Figure 8B-B’).

**Figure 8.**
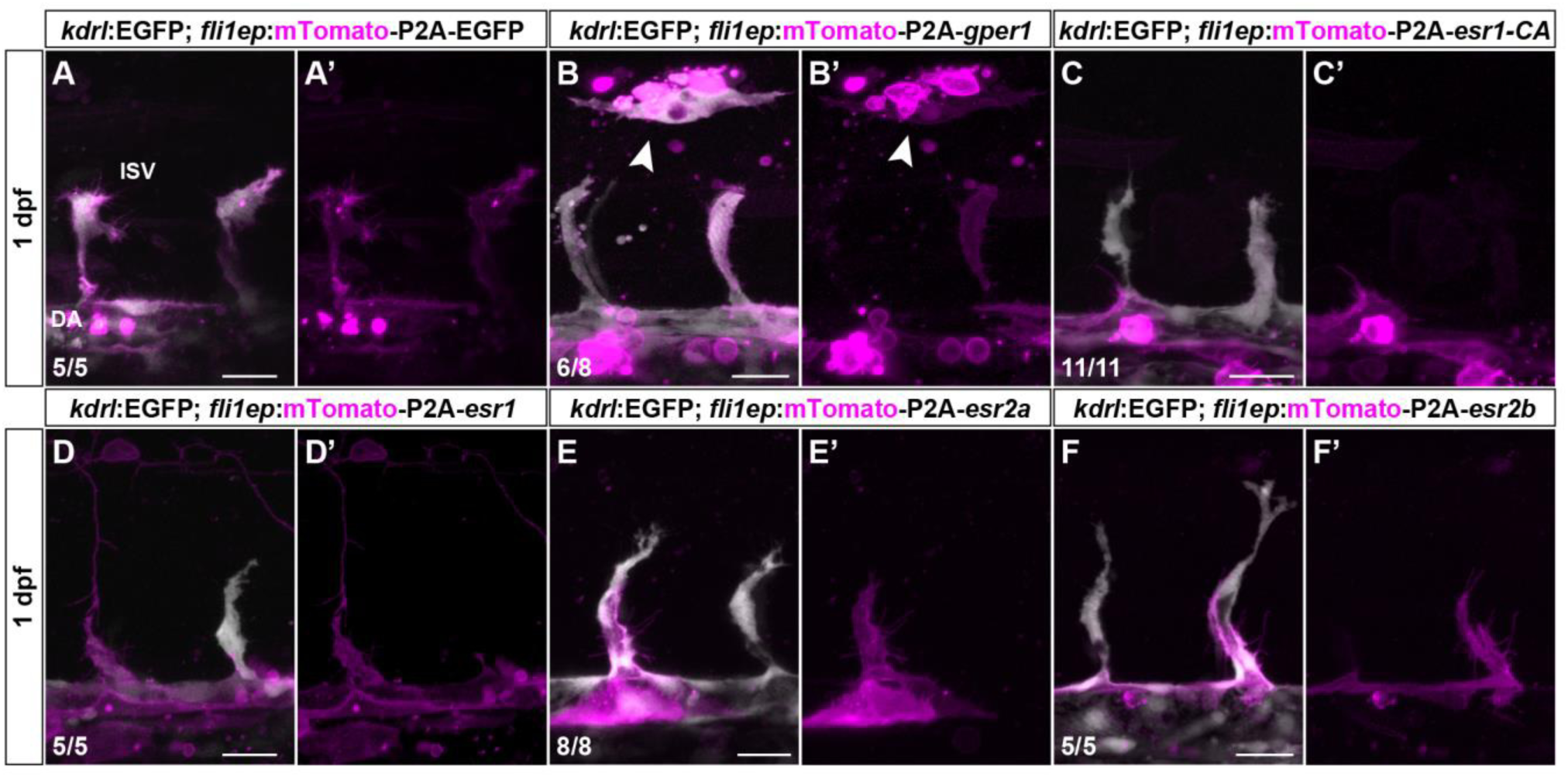
Gper1 can cell-autonomously induce EC disconnections during angiogenesis. **(A-F’)** Maximum intensity projections of endothelial mosaic overexpression of estrogen receptors. Endothelial overexpression of *gper1* induces cell-cell disconnections during developmental angiogenesis. Disconnected ECs locate to the dorsal side of the embryo (white arrowheads in (B-B’)). No overt vascular phenotypes were observed in control embryos (A-A’), or in embryos overexpressing *esr1-CA* (C-C’), wild-type *esr1* (D-D’), *esr2a* (E-E’) or *esr2b* (F-F’) in ECs. Scale bars, 25 µm. Number of embryos analyzed are shown. DA, dorsal aorta. ISV, intersegmental vessel.

We generated an *esr1* mutant (p.Y549S), recreating a mutation within the ligand-binding domain previously shown to confer constitutive activity (CA) ^37^. Injection of *fli1ep:mTomato-P2A-esr1-CA* in 1-cell stage *Tg(ERE:GFP)* embryos induced *GFP* expression in mTomato+ ECs (Supplemental Figure 6), confirming that this Esr1 mutant is also constitutively active in zebrafish. However, mosaic endothelial *esr1*-CA overexpression had no effect on angiogenesis by 24 hpf (Figure 8C-C’). Similarly, mosaic endothelial overexpression of wild-type *esr1, esr2a*, or *esr2b* had no impact on vascular morphogenesis at 24 hpf (Figure 8D-F’).

Altogether, these results indicate that Gper1 can act cell-autonomously to promote EC migration and cell-cell disconnections. Furthermore, it suggests that Esr1 acts most prominently in a paracrine fashion, possibly by negatively regulating Hh/Vegf/Notch signaling.

## DISCUSSION

In extra-ovarian tissues, estrogens are synthesized from circulating androgens and sulfated estrogens by the aromatase enzyme or the sulfatase (STS) and 17β-hydroxysteroid dehydrogenase type 1 (HSD17B1) enzymes, respectively. Increased expression of these enzymes leads to estrogen excess ^38,39^. In addition to genetic sources of estrogen excess, endocrine disruptors with estrogenic activity have been described ^24^. In contrast to beneficial proangiogenic effects of physiologic estrogens, epidemiologic and *in vivo* studies suggest that excess estrogens may induce vascular defects ^21–23^. Yet, how excess estrogen signaling impacts vascular morphogenesis remains elusive. We found that excess estrogens during early zebrafish development lead to vascular malformations due to decreased Hh/Vegfaa/Notch signaling, and increased PI3K and Rho GTPases activity. Esr1 and Gper1 are the main effectors of excess estrogens. Gper1 can cell-autonomously promote endothelial migration and cell-cell disconnections, whilst Esr1 acts in a cell non-autonomous fashion on endothelial cell behavior.

ESR1 promotes *VEGFA* expression upon binding to EREs within the proximal promoter region. Increased *Vegfa* expression induced by physiologic E2 concentrations promotes capillary formation *in vivo* and *in vitro* ^7–9^. In contrast to physiologic conditions, we show that excess estrogens impair *vegfaa* expression by negatively regulating Hh signaling. E2-dependent Vegfaa downregulation has been reported at earlier stages of vascular development, however, in contrast to our study, the authors concluded it was Hh signaling independent ^51^. Differences on tissue and developmental stage-specific estrogenic responses along with dose-dependent responses to E2 might explain the differences between our findings and previous studies. Opposing effects of estrogen signaling on Hh pathway activation have been reported in breast and gastric cancer ^40–42^, whilst neonatal estrogen exposure negatively regulates Hh signaling in the prostate ^43^. Our data confirms excess estrogen signaling as a negative regulator of Hh signaling during development, and together with previous studies, it suggests that the effects of estrogens on Hh signaling might be context and dose-dependent.

Expression of the Notch ligand gene *dll4* was also reduced in E2-treated embryos. Loss of the Notch effector genes *hey2* and *efnb2a* results in arterial-venous shunts due to increased PI3K-dependent ventral sprouting from the DA primordium ^5,6,44^. Excess estrogens did not affect *hey2* and *efnb2a* expression by 30 hpf, shortly before the onset of the DA phenotype. However, E2 treatment exacerbated the DA phenotype in *dll4* morphants, suggesting that Notch signaling is downregulated by excess estrogens. Furthermore, PI3K inhibition partially rescued the axial vessel segregation defects. These findings indicate that excess estrogens impair Notch-dependent arterial-venous segregation. However, reduced *dll4* expression did not appear to have a role in the etiology of the E2-induced ISV phenotype. During angiogenesis, DLL4 activates Notch signaling in receiver cells, which blocks EC response to VEGFA and hence prevents hypersprouting and branching ^1^. E2-treatments did not recapitulate the hyperbranching phenotype induced by deficient Notch signaling. Furthermore, injections of a *dll4* MO had no effects on the E2-induced ISV phenotype. These results are most likely explained by the reduced *vegfaa* levels in E2-treated embryos.

In addition, our study shows that *kdrl* haploinsufficiency protects against the E2-induced vascular phenotypes. Furthermore, we show that E2 treatment promote EC migration during CCV formation, a Vegfaa-independent process. Activation of PI3K/AKT signaling, downstream of integrins and receptor tyrosine kinases, initiates intracellular responses leading to the activation of small Rho GTPases. Polarized activation of Cdc42, Rac1 and RhoA is required for directional cell migration ^3,4^. In agreement with previous studies ^14,17,45–48^, our pharmacological data shows excess estrogen signaling promotes an increased activation of PI3K and Rho GTPase signaling. Importantly, it also indicates that hyperactivation of these pathways induces cell-cell disconnections within the intersegmental and axial vessels. In addition to cell migration, Rho GTPases also participate in lumen formation. Overexpression of constitutively active and dominant negative Cdc42 and Rac1 disrupts lumen formation *in vitro* ^49^. Interestingly, the onset of the E2-induced cell-cell disconnections coincides with lumen formation. Our data suggest that excess estrogens, by promoting Rho GTPases activity, may not only affect EC migration but also the maturation of cell-cell contacts and lumen expansion.

Extensive research on ESR1 and ESR2 genomic signaling indicate that these receptors have unique, overlapping and opposing actions in a ligand, cellular and genomic context dependent fashion ^50^. Our study shows that excess estrogens predominantly signal via Esr1 and Gper1, albeit a cooperative synergism with Esr2a and Esr2b may contribute to the etiology of the vascular defects. In addition to promoting *Vegfa* expression, ESR1 and GPER1 signaling from the plasma membrane promotes EC migration via activation of PI3K/AKT signaling and downstream effectors in the endothelium ^14,17,45–48^. We show that endothelial Gper1 overexpression is sufficient to induce cell-cell disconnections and excessive migration, demonstrating for the first time a cell-autonomous role for Gper1 signaling in the etiology of vascular malformations. Mosaic endothelial overexpression of wild-type or CA ^37^ nuclear estrogen receptors had no effects on EC sprouting or migration during developmental angiogenesis. These results strongly indicate that nuclear estrogen receptor signaling is the main driver of the non-endothelial response to excess estrogens leading to vascular defects.

In summary, we established a new zebrafish model for studies into dysregulated estrogen signaling during development. Using this model, we uncovered complex endothelial and extra-endothelial responses contributing to the vascular defects due to developmental estrogen excess. Importantly, our study brings new insights into the role of dysregulated estrogen signaling on the developmental programming of vascular disease. Furthermore, the identified estrogen-modulated pathways play key roles during the revascularization of ischemic tissues as well as tumor vascularization and invasiveness. Future studies using specific disease models will provide a more comprehensive understanding of how excess estrogens modulate pathologic angiogenesis, and may help to identify new therapeutic targets for improved personalized medicine.

## ACKNOWLEDGEMENTS

We thank Michele Marass and Andrea Rossi for sharing *flt1, mflt1* and *vegfaa* mutants, Radhan Ramadass and Jenny Pestel for help with imaging and image analyses, Rubén Marín-Juez, Michelle Collins and Sébastien Gauvrit for comments on the manuscript, and other members of the Stainier lab for sharing reagents and discussions. This work was supported by a postdoctoral fellowship from the European Society for Endocrinology to SP and funds from the Max Planck Society to DYRS.

---

**Movie 1A.** ISV formation and stabilization in control embryos. Related to Figure 1D. Time-lapse images of the posterior region of the trunk vasculature were acquired every 20 min between 29 and 44 hpf. Negative LIFEACT-GFP (black) and NLS-mCherry (red) expression are shown. ISVs extend and reach the dorsal side of the embryo to start forming the DLAV. Once they have migrated dorsally, ISVs are stabilized and lumen formation starts. Scale bar, 20 µm.

**Movie 1B-C.** ISV defects in E2-treated embryos due to EC disconnections. Related to figure 1D’. Time-lapse images of the posterior region of the trunk vasculature were acquired every 20 min between 29 and 44 hpf. Negative LIFEACT-GFP (black) and NLS-mCherry (red) expression are shown. The extension of the ISVs and formation of the DLAV are delayed in E2-treated embryos. Fully extended ISVs undergo stenosis and disconnect from the dorsal aorta (DA) or the DLAV. Scale bar, 20 µm.

**Movie 2A.** Extension of the DA lumen in control embryos. Related to Figure 2C. Time-lapse images of the posterior region of the trunk vasculature were acquired every 20 minutes between 28 and 43 hpf. Negative LIFEACT-GFP expression in ECs is shown. The DA is pseudocolored in red. Two representative cells are rendered in green to highlight cell shape changes during lumen extension within the DA. Scale bar, 20 µm.

**Movie 2B.** Premature truncation of the lumenized DA in E2-treated embryos. Related to Figure 2C’. Time-lapse confocal images of the posterior region of the trunk vasculature of embryos treated with E2 were acquired between 30 and 43 hpf. Negative LIFEACT-GFP expression in ECs is shown. The DA is pseudocolored in red. Two cells detaching from anterior neighbors and migrating to the posterior region of the DA are rendered in yellow and blue. Detachment of these cells from the DA leads to a premature truncation of the circulatory loop. Scale bar, 20 µm.

**Movie 3A.** EC migration in the forming CCV in control embryos. Related to Figure 5 (A-B). Time-lapse confocal images of the forming CCV in control *LIFEACT-GFP;NLS-mCherry* embryos between 32-36 hpf. Images were acquired every 15 min. LIFEACT-GFP expression is rendered in white and NLS-mCherry in red. Migration tracks of individual ECs are shown (white lines). Scale bar, 50 µm.

**Movie 3B.** EC migration in the forming CCV in E2-treated embryos. Related to Figure 5A’-B’. Time-lapse confocal images of the forming CCV in E2-treated *LIFEACT-GFP;NLS-mCherry* embryos between 32-36 hpf. Images were acquired every 15 min. LIFEACT-GFP expression is rendered in white and NLS-mCherry in red. Migration tracks of individual ECs are shown (white lines). EC nuclei and migration tracks of representative cells with decreased migration directionality are rendered in blue. Scale bar, 50 µm.

